# Elexacaftor/tezacaftor/ivacaftor treatment is associated with epigenetic age deceleration in cystic fibrosis

**DOI:** 10.64898/2026.09.13.751273

**Authors:** Josh P. Dyce, Ana I Hernandez Cordero, Michael S Kobor, Julie MacIsaac, Amrit Singh, Janice M Leung, Bradley S. Quon

**Author notes:** Corresponding Author: Bradley Quon. Co-authorship (Josh P. Dyce, Ana I Hernandez Cordero).

## Abstract

**Background:** Cystic fibrosis (CF) is associated with chronic inflammation and progressive multisystem disease, which may contribute to accelerated biological aging. Elexacaftor/tezacaftor/ivacaftor (ETI) substantially improves clinical outcomes in people with CF (pwCF), but its effects on epigenetic aging and their relationship with inflammation remain poorly understood.

**Methods:** We performed a pilot longitudinal study of 8 modulator-naïve pwCF. Blood DNA methylation was profiled before ETI initiation and after 1 year of treatment. DNAmGrimAge2 age acceleration was calculated adjusting for chronological age and blood cell composition. Associations with inflammatory biomarkers and clinical measures were assessed using repeated measures correlation.

**Results:** DNAmGrimAge2 was strongly correlated with chronological age at baseline and follow-up. DNAmGrimAge2 age acceleration decreased after ETI treatment, from a median of 1.92 years before treatment to ™1.70 years after 1 year (p=0.008). Greater age acceleration was associated with higher IL-6, IL-1β, and calprotectin concentrations. Higher FEV1pp was associated with lower age acceleration, whereas higher sweat chloride was associated with greater age acceleration.

**Conclusions:** DNAmGrimAge2 age acceleration decreased following ETI initiation and was associated with markers of innate and neutrophilic inflammation and CF disease activity. These findings suggest that epigenetic aging in CF may be dynamic and responsive to improvements in CFTR function and inflammatory disease burden. Larger longitudinal studies are needed to validate these findings and determine their relationship with long-term clinical outcomes.

## Introduction

Cystic fibrosis (CF) is characterized by chronic inflammation and progressive multisystem disease, processes that may promote accelerated biological aging. Chronic inflammation has been proposed as an important driver of “inflammaging” in CF ^1^, but whether restoration of CFTR function modifies biological aging is poorly understood. Epigenetic clocks derived from DNA methylation provide quantitative biomarkers of biological aging, with DNAmGrimAge2 incorporating methylation signatures associated with age-related morbidity and mortality. A recent study reported an association between epigenetic age and lung function in people with CF (pwCF) treated with elexacaftor/tezacaftor/ivacaftor (ETI)^2^; however, the relationship between ETI-associated changes in epigenetic aging and inflammation has not been investigated. We therefore evaluated longitudinal changes in DNAmGrimAge2 age acceleration from modulator-naïve baseline to one year after ETI initiation and explored whether epigenetic aging was associated with circulating inflammatory biomarkers and clinical measures of CF disease activity.

## Methods

### Study Cohort

This pilot study included eight modulator-naïve pwCF enrolled in the CAN-IMPACT study^3^, an observational study investigating biological changes following ETI initiation. Biobanked peripheral blood collected immediately before ETI initiation and at one-year follow-up were used for DNA methylation profiling and measurement of inflammatory biomarkers. Participants provided consent for the use of their samples for future research (UBC-PHC-REB H21-01707).

### DNA Methylation Assay

DNA was extracted using the DNeasy Blood and Tissue Kit (Qiagen, Hilden, Germany) and bisulfite converted using the EZ DNA Methylation Kit (Zymo Research, Irvine, California). DNA methylation was profiled using the Infinium MethylationEPIC v2.0 array (∼ 930k CpG sites) with samples randomized within chips to minimize batch effects. CpG probes were excluded if they had poor detection quality (*P* > 1 × 10^−10^), mapped to non-CpG sites, overlapped single nucleotide polymorphisms, or were cross-hybridizing.

Background correction, normalization, and batch correction were performed using Noob^4^, Beta-Mixture Quantile Normalization^5^, and ComBat^6^, respectively, according to our previously established pipeline^7^. DNAmGrimAge2 was calculated using the Clock Foundation webtool^8,9^. We selected DNAmGrimAge2 because GrimAge-based clocks have demonstrated associations with morbidity and mortality and because our previous work found DNAmGrimAge to best approximate airway epigenetic age and predict clinical outcomes in chronic lung disease^10,11^.

Estimated blood-cell proportions were generated using the Houseman deconvolution method^12^. DNAmGrimAge2 age acceleration was calculated as the residual from regression of DNAmGrimAge2 on chronological age while adjusting for the first principal component derived from estimated blood-cell proportions. Positive residuals therefore represent an epigenetic age older than expected for chronological age.

### Inflammatory Biomarker Assays

Plasma inflammatory biomarkers were measured using Meso Scale Discovery multiplex immunoassays. The panel encompassed biomarkers representing several inflammatory pathways relevant to CF, including neutrophilic inflammation (calprotectin), innate immune activation (IL-1β, IL-6, TNF-α), type 2 immune responses (IL-4, IL-5, IL-13), and Th17-associated inflammation (IL-17A), together with MIP-3α, a chemokine involved in leukocyte recruitment. C-reactive protein (CRP) was measured as an acute-phase inflammatory protein. Complete blood counts and leukocyte differentials were also evaluated.

### Statistical Analysis

Pre- and post-ETI age acceleration residuals were compared using paired Wilcoxon signed-rank tests. Repeated-measures correlation was used to assess within-participant relationships between age acceleration and the inflammatory/immune biomarkers across visits. Biomarker *P*-values were corrected using the Benjamini-Hochberg false discovery rate (FDR), with FDR-adjusted *P*<0.05 considered significant. Associations with prespecified clinical measures (FEV1pp and sweat chloride) were evaluated using repeated-measures correlation without multiple-comparison adjustment. Analyses were performed in R v4.4.2.

## Results

### Demographics and Clinical Characteristics

Participants had a median age of 25.4 years (IQR 22.1–28.0), and two (25%) were female. Five were homozygous for F508del and three were F508del heterozygous. No individuals had a prior history of CFTR modulator use. Baseline median FEV1pp was 64.5% (IQR 53.8–75.0) and sweat chloride was 98.0 mmol/L (IQR 96.0–103.8). After one year of ETI, FEV1pp increased to 90.0% (IQR 75.2–98.8; *P*=0.014) and sweat chloride decreased to 41.0 mmol/L (IQR 37.8–67.8; *P*=0.008). The most commonly isolated organisms were methicillin-susceptible *Staphylococcus aureus* (MSSA) in 50% of patients and *Pseudomonas aeruginosa* in 25% of patients.

### Epigenetic Age Correlations

DNAmGrimAge2 correlated strongly with chronological age both before (R=0.89, *P*=0.003) and after ETI (R=0.85, *P*=0.007) (**Figure 1A**). DNAmGrimAge2 age acceleration decreased significantly following ETI (*P*=0.008; **Figure 1B**). Before ETI, participants had a median age acceleration of 1.92 years (IQR 0.12–2.65), whereas after one year of ETI, DNAmGrimAge2 was a median 1.70 years younger than expected for chronological age (IQR ™1.97 to ™1.17).

**Figure 1.**
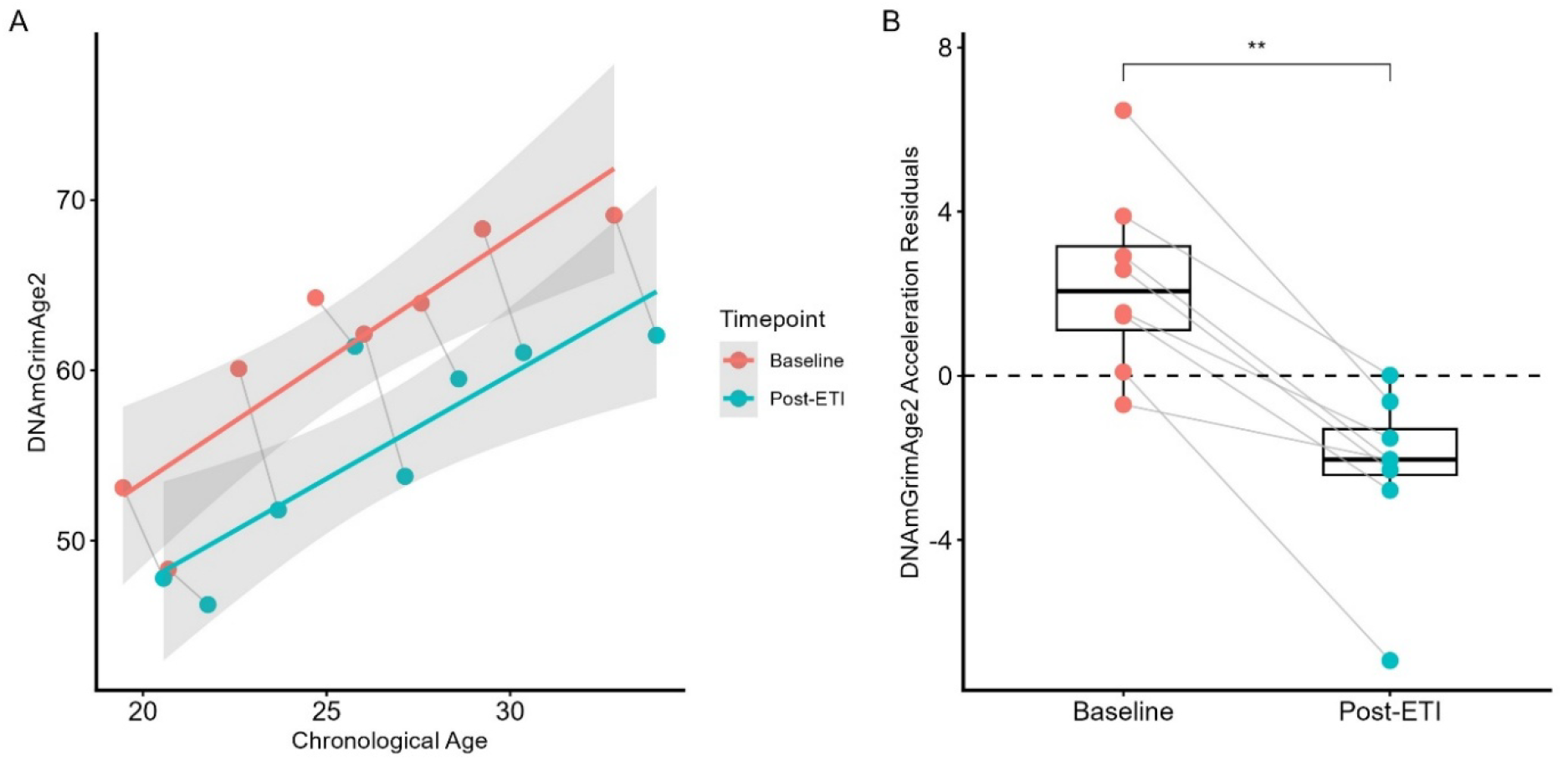
Effect of ETI treatment on DNAmGrimAge2 age acceleration. (A) Associations between chronological age and DNAmGrimAge2-predicted age at baseline and post-ETI. Lines represent separate linear regression fits for each time point with 95% CIs; estimates are not adjusted for blood cell composition. (B) Blood cell composition-adjusted DNAmGrimAge2 age acceleration residuals at baseline and post-ETI. Paired values were compared using the Wilcoxon signed-rank test (** p ≤ 0.01).

To explore whether this change in biological aging tracked with inflammation, we examined within-participant associations between DNAmGrimAge2 age acceleration and circulating biomarkers (**Figure 2**). Higher IL-6, IL-1β, and calprotectin were significantly associated with greater age acceleration after FDR correction (FDR adjusted p < 0.05). Several additional markers, including CRP and total white blood cell and lymphocyte counts, showed positive but non-significant associations.

**Figure 2.**
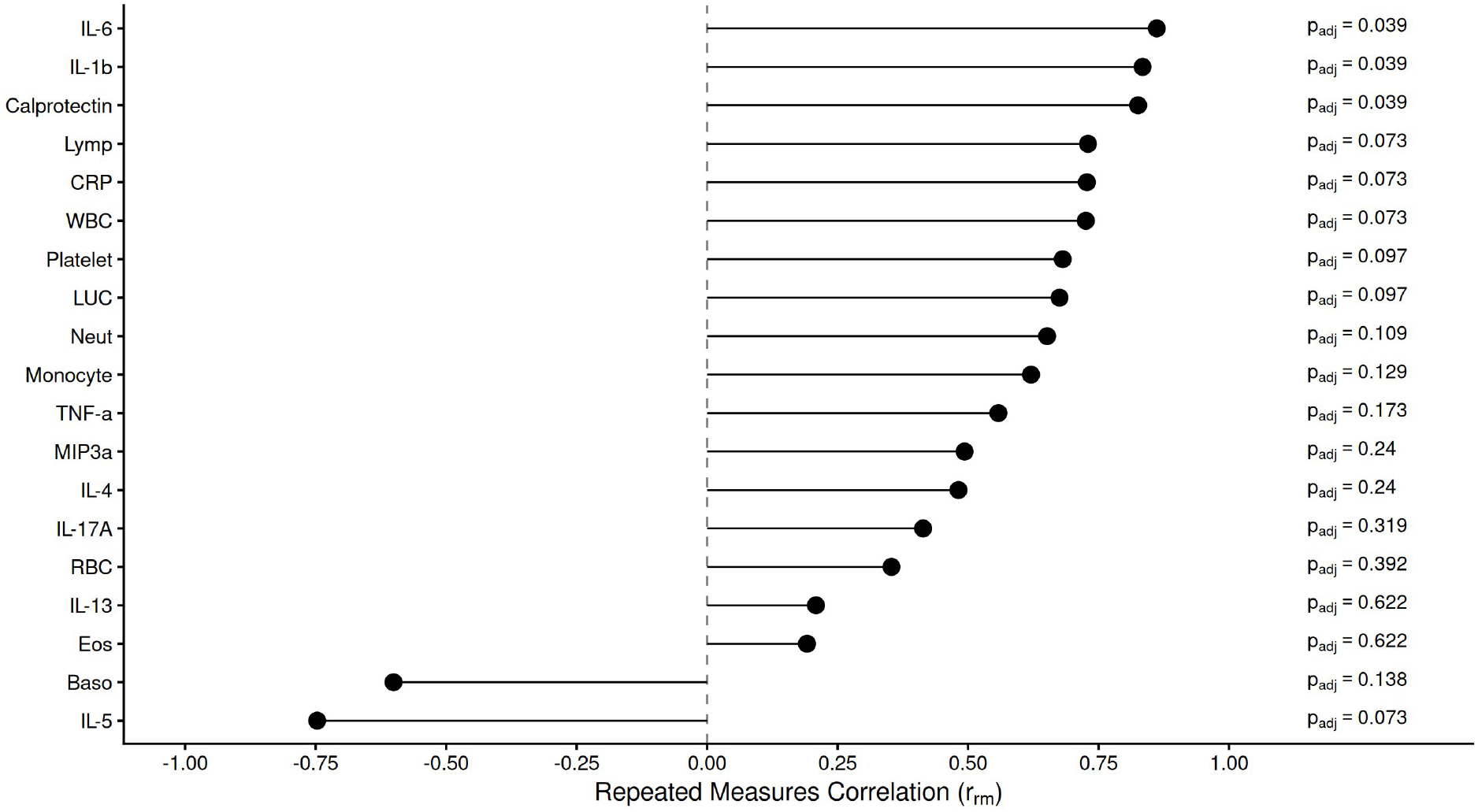
Association between biomarkers and DNAmGrimAge2 age acceleration. Repeated measures correlation used to assess within-participant associations between biomarker levels and DNAmGrimAge2 age acceleration across visits. Positive r_rm_ indicate that higher biomarker levels were associated with greater age acceleration, whereas negative values indicate the opposite relationship. P-values testing whether r_rm_ differed from zero were adjusted for multiple comparisons using the Benjamini-Hochberg false discovery rate procedure.¶

DNAmGrimAge2 age acceleration was also associated with clinical measures of CF disease activity (**Figure 3**). Higher FEV1pp was associated with lower epigenetic age acceleration, whereas higher sweat chloride concentration was associated with greater epigenetic age acceleration.

**Figure 3.**
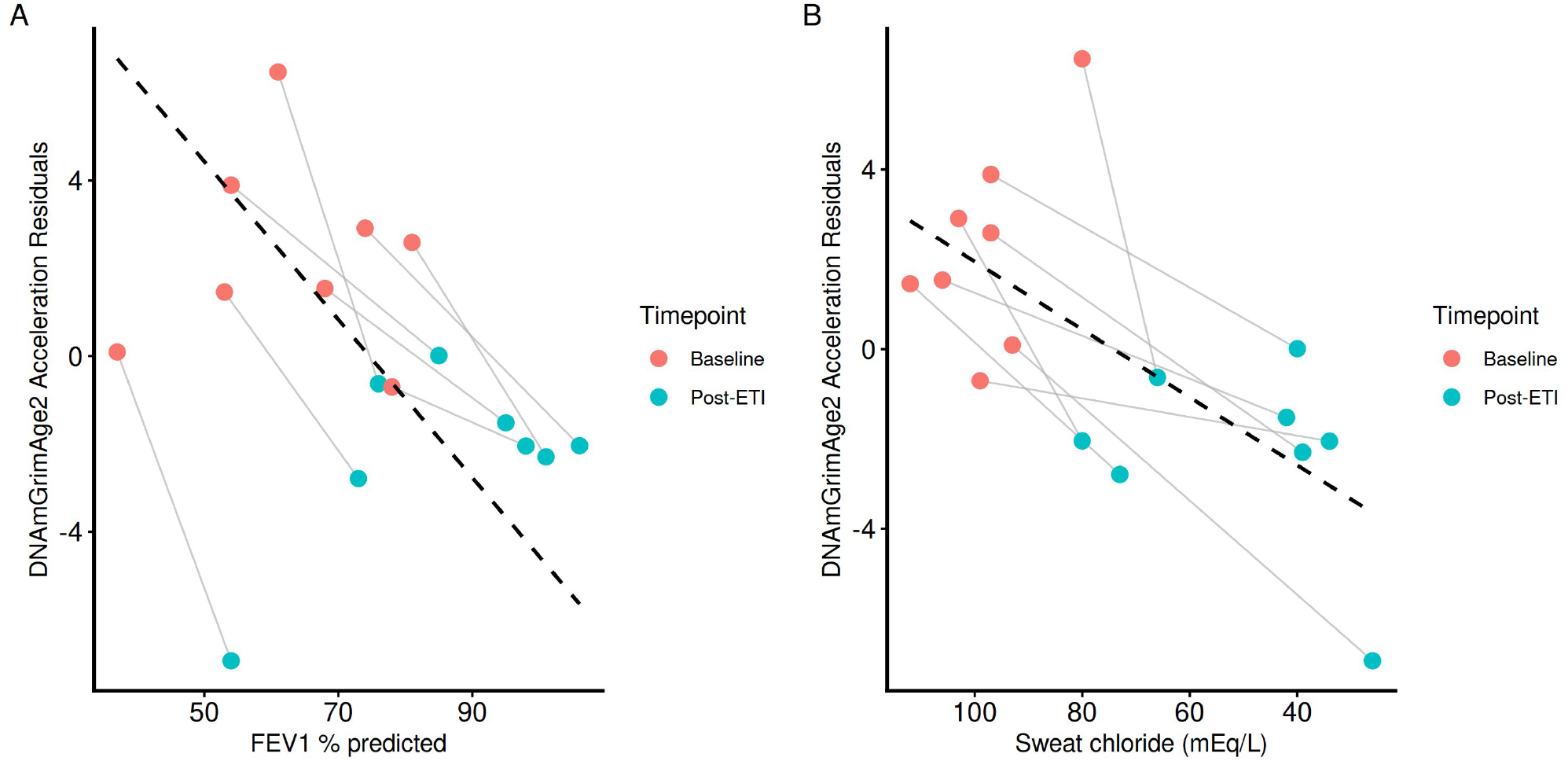
Association between clinical measures and DNAmGrimAge2 age acceleration. Repeated measures correlation was used to assess within-participant associations of: (A) FEV1 % predicted and (B) sweat chloride concentration with DNAmGrimAge2 age acceleration. Higher FEV1 % predicted was associated with lower age acceleration (r_rm_ = -0.86 *p* = 0.003), whereas higher sweat chloride was associated with greater age acceleration (r_rm_ = 0.80, *p* = 0.009). Points represent individual measurements, with lines connecting repeated measurements from the same participant. Dashed lines represent the common within-participant regression slopes estimated by the repeated measures correlation model.

## Discussion

In this longitudinal pilot study of modulator-naïve pwCF, DNAmGrimAge2 age acceleration decreased significantly during the first year of ETI treatment. Importantly, greater age acceleration tracked within individuals with higher circulating IL-6, IL-1β, and calprotectin, providing preliminary evidence linking epigenetic aging in CF with systemic innate and neutrophilic inflammation. Together, these findings suggest that the profound clinical effects of CFTR modulation may extend to biological processes associated with aging and that attenuation of chronic inflammation may accompany these changes.

Our findings extend the recent observations of Castaldo et al. who reported an inverse correlation between epigenetic age and FEV1pp in 52 adults with CF and changes in epigenetic age following one year of ETI^2^. Our study differs in several important respects. We studied participants from a modulator-naïve baseline, used DNAmGrimAge2 rather than the Horvath Skin & Blood clock^13^ to better approximate lung aging, calculated age acceleration using age- and blood-cell-composition-adjusted residuals rather than an epigenetic-to-chronological-age ratio, and evaluated inflammatory correlates of epigenetic aging. The association between greater age acceleration and lower FEV1pp is consistent with their findings, whereas our association with sweat chloride was not observed in their cohort.

The relationships with IL-6, IL-1β, and calprotectin are particularly notable given the proposed role of chronic inflammation in accelerated aging in CF. Persistent neutrophilic and innate immune activation can promote oxidative stress, cellular injury, and senescence, providing a plausible biological link between CF disease activity and epigenetic aging. The simultaneous improvement in CFTR function, lung function, systemic inflammation, and DNAmGrimAge2 following ETI raises the possibility that restoration of CFTR function modifies an inflammation-associated biological aging phenotype. However, these correlated changes cannot establish whether reduced inflammation mediates the observed change in epigenetic aging.

The small sample size is the principal limitation of this pilot study and precluded multivariable modeling to determine whether inflammatory biomarkers, lung function, and CFTR function are independently associated with DNAmGrimAge2. The observational pre/post design also cannot distinguish an ETI-specific effect from other time-varying factors, and the findings require replication in larger cohorts. Nevertheless, the longitudinal assessment of modulator-naïve individuals provides preliminary evidence that epigenetic aging in CF may be dynamic rather than solely reflecting accumulated irreversible disease burden. Larger longitudinal studies should determine whether ETI-associated epigenetic age deceleration is sustained and whether residual or progressive epigenetic age acceleration identifies individuals with persistent inflammation or increased risk of future CF-related complications.

